# Niche conservatism in the Legume Amherstieae tribe: Insights from the tropical Berlinia and Brownea clades

**DOI:** 10.1101/2024.09.12.612774

**Authors:** Ingrid C. Romero, Surangi W. Punyasena

## Abstract

The concept of niche conservatism describes the tendency of organisms to retain ecological traits through time and space. Reviewing this concept in different groups of angiosperms is critical to understanding what factors drove their expansion and geographic distribution, as well as assessing how, in taxonomic levels higher than species, ecological traits have remained relatively constant through time and space. Studying niche conservatism can also help us understand how the distribution of clades may be affected by climate change. Niche conservatism has been observed in many clades of legumes. Amherstieae, the largest tribe of the Detarioideae subfamily, has a geographic distribution, evolutionary history, and phylogeny that makes it a good candidate for evaluating patterns in niche conservatism. We analyzed the distribution of two Amherstieae suprageneric clades, the Berlinia and Brownea clades. The former is endemic to sub-Saharan Africa and the latter is restricted to the Neotropics. We used the geographic distributions of each clade to define their G-space (geographic space) and extracted 19 climatic variables to define the E-space (environmental space) of each clade. We used two tests to evaluate the similarity in the climatic niche of both clades, the niche overlap test (NOT) to estimate similarities between the occupied E-spaces (realized niche space) and the niche divergence test (NDT) to assess the similarity of the environmental niche relative to the accessible analogous E-space (potential niche space) of each biogeographic region. Our results suggest that the Brownea clade are descendants of a climatic subset of the Berlinia clade preferring less variable temperature and higher precipitation levels, and that the dry-adapted subset of Berlinia may represent a more recent evolutionary expansion.

## INTRODUCTION

The concept of niche conservatism has been reviewed extensively at the species level, but few studies have evaluated its occurrence at higher taxonomic levels (Hadly et al., 2009; Wiens & Graham, 2005). At the species level, the current geographic ranges occupied by species are considered a proxy for their realized or occupied niche (Hutchinson, 1958; Hadly et al., 2009) but they do not necessarily indicate the full range of environments species can potentially inhabit – their fundamental niche (Brown & Carnaval, 2019; Jackson & Overpeck, 2000; Wiens & Graham, 2005). Jackson and Overpeck (2000) introduced the concept of “potential niche” as a subset of the fundamental niche – the intersection of the fundamental niche with the abiotic and biotic conditions at a specific point in time. A change in a species geographic range or distribution can be a response to environmental change, representing one possible outcome of the fundamental niche (Brown & Carnaval, 2019; Hadly et al., 2009; Jackson & Overpeck, 2000). However, the fundamental niche may also be constrained at higher taxonomic levels due to intrinsic life history and developmental traits (Crisp et al., 2009; Hadly et al., 2009; Wiens et al., 2010; Wiens & Graham, 2005).

Measuring the degree of niche conservatism in taxonomic clades at a rank higher than individual species allows us to understand broad-scale patterns of biodiversity, as well as predict clade-specific responses to climate change (Crisp et al., 2009; Hadly et al., 2009; Qian & Ricklefs, 2004). For example, species with narrow, conserved thermal tolerances may be at increased risk of warming temperatures (Crisp et al., 2009; Wiens & Graham, 2005). Most legumes adapted to tropical dry environments, such as seasonally dry tropical forests and savannas, tend to conserve these environmental niches (Oliveira-Filho et al., 2013; Pennington & Lavin, 2016), while legumes with ancestors adapted to tropical wet environments, such as Detarioideae (de la Estrella et al., 2020, 2017; Murphy et al., 2018; Schley et al., 2018) display more environmental diversity, and include dry-adapted species (de la Estrella et al., 2020, 2017).

Detarioideae, a monophyletic subfamily of Leguminosae, is among the early-branching clades in legume evolution (de la Estrella et al., 2018). The largest tropical tribe within Detarioideae, Amherstieae, is primarily adapted to wet tropical environments (Dataset S1). Originating in Africa, this tribe’s diversity is mainly concentrated in the Paleotropics (Figs. 1, de la Estrella et al., 2017, 2020; LPWG, 2017; Romero et al., 2020; Schley et al., 2018). Amherstieae consist predominantly of tree species distributed in wet lowland forests, along rivers, on rock slopes, and in sandy and loamy soils, with only a few species distributed in drier habitats (Dataset S1, Bruneau et al., 2008; de la Estrella et al., 2017; Mackinder & Pennington, 2011). Despite similar ecological characteristics across its genera, only three are pantropical and 93% are restricted to specific regions such as Asia, Africa, or tropical America (Table S1, de la Estrella et al., 2017; de la Estrella et al., 2020; Murphy et al., 2018; Schley et al., 2018). This geographic specialization highlights the diverse adaptation strategies within the tribe.

**Figure 1.**
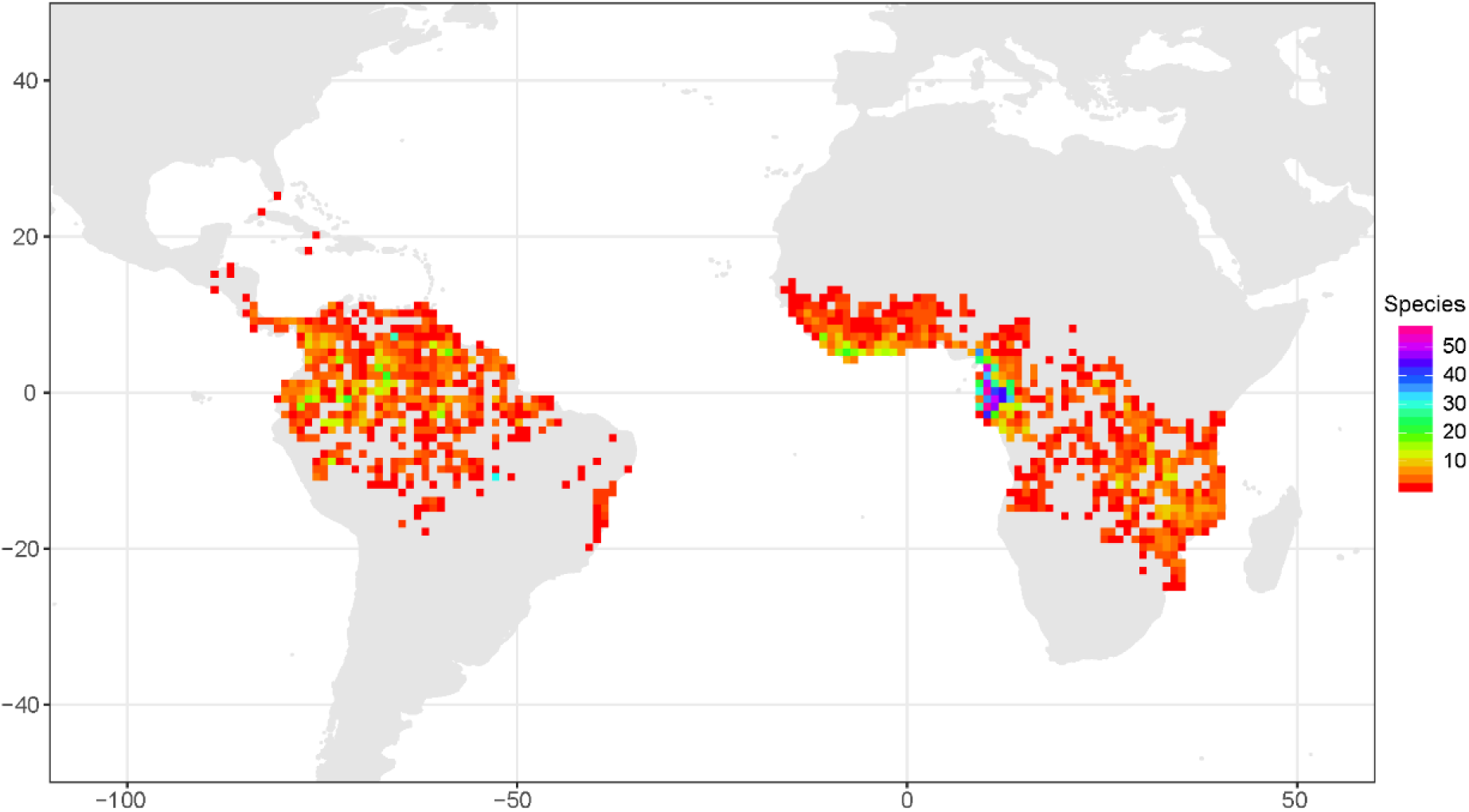
Heatmap of species richness of the Berlinia and Brownea clades, Amherstieae tribe. The color scale indicates the number of species. Grid size: ∼55.5 Km.

To assess the strength of climatic niche conservation in Amherstieae across geographic regions, we compared the geographic and climatic distributions of the African Berlinia clade (16 genera, ∼179 species, Fig. S1) and the Neotropical Brownea clade (9 genera, ∼113 species, Fig. S2). Both supra-generic clades are monophyletic (Schley et al., 2018; de la Estrella et al., 2020). Combined they include almost 50% of Amherstieae genera and are endemic to two distinct biogeographic regions (Table S1, de la Estrella et al., 2020; Schley et al., 2018). They are composed of species that are dominant ecological components of tropical forests (Bruneau et al., 2008, 2014; de la Estrella et al., 2018; Murphy et al., 2018; Ter Steege et al., 2013), or economically important as timber sources (de la Estrella et al., 2018; Mackinder & Pennington, 2011; Wieringa, 1999). Phylogenetically, Berlinia is the closest African relative of the Neotropical Brownea clade (Mackinder & Pennington, 2011; Murphy et al., 2018; Schley et al., 2018). The African clade originated early in the Cenozoic, >60-40 million years ago (Ma), but most genera are relatively young, originating in the Miocene and Pleistocene (12-1 Ma, de la Estrella et al., 2020; Demenou et al., 2020). The Brownea clade, which contains all the Neotropical genera of the tribe, originated during the Eocene (∼34 Ma, de la Estrella et al., 2017; Schley et al., 2018), with most of the diversification occurring in Amazonia during the Middle Miocene (12-7 Ma, Schley et al., 2018).

Analyzing the environmental niches of the Berlinia and Brownea clades gives us insights into the climatic factors that drive the environmental distributions of Amherstieae and may allow us to predict how these species will be impacted by future climate. If there is strong niche conservatism in Amherstieae, we expect to see similarities in the environmental space occupied by the Berlinia and Brownea clades, despite their large geographic and long evolutionary separation. If there is little niche conservatism, we expect to see a variation in the environmental space occupied by the two clades that reflect climatic differences between the Neotropics and the Paleotropics. Similarities and differences in environmental niches are a result of the evolutionary history of these clades.

## METHODS

### Geographic data of the Amherstieae clades

The Berlinia and Brownea clades are considered sister taxa. They diverged in the early Cenozoic and have since diversified in separate continents (Bruneau et al., 2008; de la Estrella et al., 2017; 2018; 2020; Schley et al., 2018). The Berlinia clade is distributed across tropical Africa (de la Estrella et al., 2020), Figs. 1, S1), while the Brownea clade is widely distributed in Central America and northern South America (Schley et al., 2018, Figs. 1, S2).

We compiled geographical generic-level occurrences of the Berlinia and Brownea clades from the Global Biodiversity Information Facility (GBIF) (Dataset S2, GBIFa; GBIFb; GBIFc; GBIFd, 2020). We reviewed and cleaned the occurrences using R and R Studio 3.5.3 packages janitor (Firke et al., 2020) and CoordinateCleaner (Zizka et al., 2019). We excluded records from GBIF indicated as: 1) fossils; 2) cultivated or introduced specimens; 3) did not have coordinates or the coordinates were zero; and 4) had invalid geographic coordinates and/or an invalid country. Our final dataset contained a total of 25,441 specimens, corresponding to 25 genera and ∼326 species (Dataset S2, Figs. S1, S2).

We transformed our point occurrences into a raster grid of 55.5 km x 55.5 km cells, in which the value of each cell corresponded to the number of species found within (Topel et al., 2017). The result was a map capturing the variation of species richness within the clades in Africa and the Americas (Figure 1). For this, we used the R packages speciesgeocodeR (Topel et al., 2017) and raster (Hijmans & van Etten, 2019)

### Definition of the geographic space

We defined two types of geographic space (G-space) by latitude and longitude (Fig. S3c,d, Brown & Carnaval, 2019; Warren et al., 2010). The first was the available G-space (_av_G-space), which describes the area that could be occupied by a taxon or group of taxa in a geographic region. The second was the occupied G-space (_oc_G-space), which corresponds to the current geographic area occupied by a taxon or group of taxa. We produced two geographical rasters for both types of G-space, _av_G-space and _oc_G-space. For _av_G-space, the rasters varied in latitudinal and longitudinal ranges, for Africa (lat: -35, 35; lon: -18, 60) and for Neotropical America (lat: - 40, 40; lon: -120, -30), while for the _oc_G-space, the rasters used the corrected GBIF occurrences of each clade (Fig. S3c, d, Dataset S2).

### Definition of the environmental space

To define the environmental spaces, we extracted climatic data for each _oc_G-space and _av_G-space raster coordinate point from WorldClim version 2.0 (Table S1, Fick & Hijmans, 2017). WorldClim provides 19 bioclimatic variables derived from monthly temperature and rainfall measurements and at 2.5-minute spatial resolution (Hijmans et al., 2005). We used the R packages raster (Hijmans & van Etten, 2019) and humboldt (Brown & Carnaval, 2019). After, we defined two multidimensional environmental spaces (E-space), the available (_av_E-space) and the occupied (_oc_E-space) (Fig S3e-h, Di Cola et al., 2017; Nunes & Pearson, 2017; Qiao et al., 2015). The _av_E-space refers to the combination of climate conditions within our _av_G-space, while the _oc_E-space refers to the combination of climate conditions found within our _oc_G-spaces.

### Definition of the analogous environment, niche indexes and tests

For the identification of shared analogous environments and the niche analyses, we used the R package humboldt (Brown & Carnaval, 2019). We first converted the rasters into point data frames by using rarefaction to reduce potential spatial autocorrelation, eliminate spatial clusters by reducing the number of coordinate points per raster, and scale the rasters to a similar spatial resolution (40 km^2^) (Fig. S3a,b, Zurell et al., 2020; Broennimann et al., 2012; Brown & Carnaval, 2019; Dormann et al., 2007; Rotllan-Puig & Traveset, 2016). Second, the E-space was characterized as a two-axes principal component analysis using the environmental variables across geographic regions for each clade (Fig. S3). Third, a kernel density function was used to create a continuous E-space surface by gridding the E-space and using the PC values from the input occurrence localities or study region data to either estimate the E-space of each clade or each of their environments, respectively (Fig. S4). Fourth, a boosted regression tree model was used to reduce the 19 environmental variables and select only those that contributed to more than 10% of the variance for either clade (Fig. 2).

**Figure 2.**
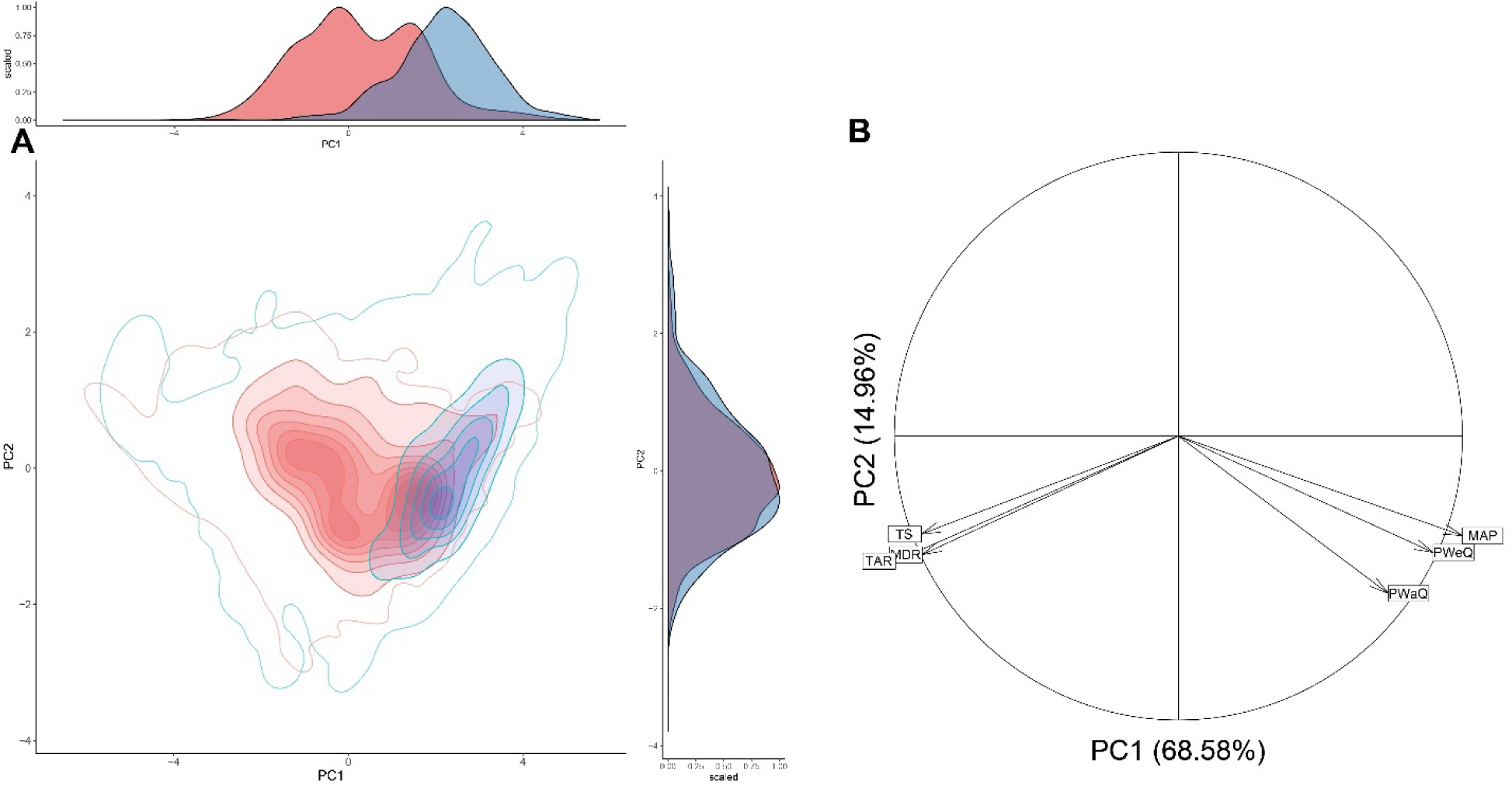
Principal component analysis of the E-space occupied and shared by the Berlinia and Brownea clades. A) Scaled kernel density plots for the first two principal components with the Berlinia clade in red and the Brownea clade in blue. The distribution of the two clades can be distinguished by PC1, but not PC2. The filled kernel density isopleths represent the space occupied by the clades. The empty kernel density isopleths represent the space available to each clade but not occupied. B) PCA vectors illustrating the direction and magnitude of only the six environmental variables that contribute the most to defining the clades’ niches in the E-spaces occupied by the Berlinia and Brownea clades. The first two principal components explain 83.54% of the environmental variation of these clades. The six variables are mean annual precipitation (MAP), precipitation of the wettest quarter (PWeQ), precipitation of the warmest quarter (PWaQ), temperature seasonality (TS), temperature annual range (TAR), and mean diurnal range (MDR). See PCA with all variables in Fig. S4.

We identified the analogous environments by comparing the E-spaces of the Brownea and Berlinia clades using different indexes and tests, such as the potential niche truncation (PNTI), niche similarity test, and niche overlap and divergence indexes. When comparing the E-spaces between the clades, the _oc_E-space represented the realized niche of each clade and the _av_E-space represented the potential niche for each biogeographic region (Brown & Carnaval, 2019; Jackson et al., 2000; Peterson, 2011).

The PNTI was used to identify whether the _av_E-space truncated the _oc_E-space. This index measures the overlap between the 5% kernel density isopleth of the E-space of the clades and the 10% kernel density isopleth of the accessible environment’s space. The PNTI value is the portion of the species’ niche perimeter that falls outside the E-space limits (Fig. 3, Brown & Carnaval, 2019). If the truncation index is high, this suggests that the realized distribution falls at the boundaries of the available environmental space and is likely an underestimate of the taxon’s fundamental niche.

**Figure 3.**
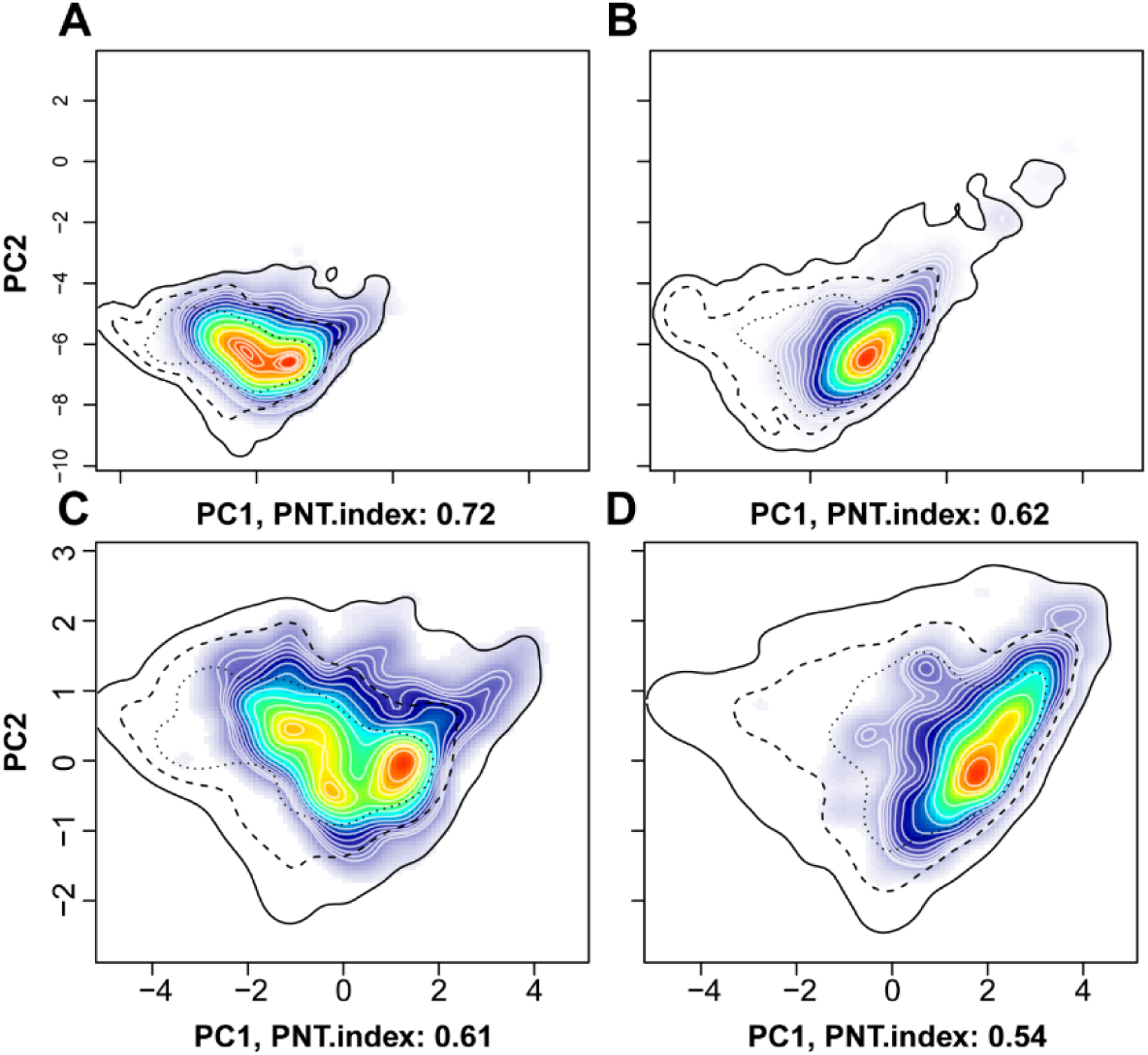
Kernel density plots representing the Potential Niche Truncation (PNT). PNT values for the Berlinia clade (A, C) and the Brownea clade (B, D). PNT values for niche overlap index (A, B) and for niche divergence index (C, D). Cooler colors (i.e. blue) represent lower densities of the E-space and warmer (i.e. red) represents higher densities. Each white line in the kernel density isopleth characterizes a density value. The black lines represent the E-space corresponding to the environment (not clades), in which: solid line=0.1, dashed line=0.5, and dotted line=0.75.

Niche similarity was measured using Schoener’s D similarity index, which equals one minus the total variation distance (a distance measure for probability distributions) between two environmental niches (Brown & Carnaval, 2019; Schoener & Gorman, 1968). Similarity ranges from 0-1, where 0 indicates niche divergence and 1 indicates niche equivalency. Additionally, we applied a left-tailed statistical test to evaluate the statistical significance (Brown & Carnaval, 2019).

We used a niche overlap test (NOT) to evaluate the similarities between the _oc_E-spaces of both clades and a niche divergence test (NDT) to test similarity in the _oc_E-spaces of both clades relative to the analogous shared climatic space (Brown & Carnaval, 2019; Di Cola et al., 2017; Warren et al., 2010). NOT and NDT calculate the overlap of the analogous and non-analogous environments occupied by both clades (Brown & Carnaval, 2019; Di Cola et al., 2017; Warren et al., 2010) and identify potential climatic adaptations that are shared between clades or unique to each clade (Alexander et al., 2015; Brown & Carnaval, 2019; Estrada et al., 2015; Garcia-Barros & Romo Benito, 2010). In the NOT, we used the entire accessible space, including shared and non-shared areas, to quantify the clades’ environments. Conversely, the NDT focused only on the shared space to quantify accessible environments, offering a detailed comparison of how each clade adapts within common climatic parameters.

For both NOT and NDT, the niche similarity and analogous environments were calculated using equivalence and background statistics (Brown & Carnaval, 2019). The equivalence statistic assesses whether two niches are equivalent based on correlative distribution models (Brown & Carnaval, 2019). This quantifies the E-space based on the niches of both clades and estimates the proportion of the accessible E-space that is shared by both clades (similarity analogous environment only - SAE). The statistics first calculate Schoener’s D similarity index, then compare it to those obtained when occurrences of the two clades are resampled by iteration (Brown & Carnaval, 2019). For each resampling, occurrences for each clade are selected and then randomly assigned to one of the clades, and, in each iteration, Schoener’s D was measured between the two reshuffled groups (Brown & Carnaval, 2019).

The background statistic aimed to evaluate the power of the equivalent test to detect differences between two groups based on available environmental conditions (Brown & Carnaval, 2019). It estimates the E-space represented within the geography of the clades, and it assesses if the clades are more different than expected given the underlying environmental differences between the regions in which they occur (Brown & Carnaval, 2019). The background statistic compares observed niche similarities between both clades to the overlap between the Berlinia clade and the random shifting of the spatial distribution of the Brownea clade in geographic space, then measures how that shift in geography changes the _oc_E-space (Brown & Carnaval, 2019).

## RESULTS

A total of 10,053 sites (4,813 for Africa and 5,240 for America) comprise the shared accessible E-space (analogous environments) between Africa and America for the Berlinia and Brownea clades (Fig S3). For the analogous E-space, the PCA indicates that 83.54% of the variance in the environmental data is explained in a 2-dimensional E-space (Fig. 2). PC1 accounted for ∼69% of the variation, indicating some differentiation between the niches of the clades, while PC2 explained 15% of the climatic variation but did not differentiate the clades (Fig. 2). The environmental variables with the highest loadings were mean diurnal temperature range (MDR), temperature seasonality (TS), annual temperature range (TAR), annual precipitation (MAP), precipitation of the wettest quarter (PWeQ), and precipitation of the warmest quarter (PWaQ) (Fig. 2, S4, S5). Along PC1, the distribution of the Berlinia clade was bimodal, and for the Brownea clade was unimodal (Figs. 2). The Brownea clade and a group of species of the Berlinia clade showed overlapping of the E-space, which were positively correlated with the precipitation variables but negatively correlated with the temperature variables (Figs. 2, S4, S5). The Berlinia clade group that did not overlap with the species of the Brownea clade was positively correlated with the temperature variables but negatively correlated with precipitation (Figs. 2, S4, S5).

In the niche overlap and niche divergence tests, the potential niche truncation index (PNTI) indicated a high potential for niche truncation in both clades (Table 1, Fig. 3). However, PNTI was slightly higher in Africa, which may be because the Berlinia clade occupies a larger percentage of the available environmental space (Fig. 3). The PNTI values indicate that the realized niches (_oc_E-space) of both the Berlinia and Brownea clades represent only a portion of their respective fundamental niches.

**Table 1.** Results from the Niche overlap (NOT) and niche divergence (NDT) tests. (*) indicates significant. NOT estimates the similarity between the occupied niches of the clades. NDT estimates the portion of analogous accessible E-space that is shared by both clades.

| Value | NOT | NDT |
| --- | --- | --- |
| PNTI Berlinia clade | 0.72 | 0.62 |
| PNTI Brownea clade | 0.61 | 0.54 |
| Niche similarity (*D*) | 0.3 | 0.59 |
| Similarity-analogous environments (SAE) | 0.67 | 0.64 |
| Equivalence p-value | 0.54 | 0.99 |
| p-value background, Brownea compared to Berlinia | 0.54 | 0.05* |
| p-value background, Berlinia compared to Brownea | 0.05* | 0.03* |

Our niche overlap analysis indicated that the occupied climatic niches of the clades are similar (NOT: D=0.3, p>0.05, SAE=0.67) (Table 1). The calculated similarity is even stronger when accounting for climatic differences in their biogeographic regions (NDT: D=0.59, p>0.05, SAE=0.64) (Table 1). The _oc_E-space of the Brownea clade was a subset of the Berlinia clade, with no portion of the occupied Brownea climatic niche unique to the clade (Fig. 2, Table 1). However, the _oc_E-space of the Berlinia clade included environmental combinations that had no overlap with the South American clade, and the shared analogous environments analysis indicated that the _oc_E-space of the Berlinia clade was significantly different from that of the Brownea clade (Fig. 2, Table 1).

## DISCUSSION

Our study explored niche conservatism in Amherstieae by comparing the environmental niches of two monophyletic clades, Berlinia and Brownea. Both clades are found in warm tropical climates (Figs. S1-2, Dataset S2). Our results showed overlapping in the _oc_E-spaces between both clades, with shared climatic preferences for less variable temperatures and adaptations to high precipitation (Fig. 2, S4). The Neotropical Brownea clade is an environmental subset of the more climatically diverse Berlinia clade. This suggests that the Neotropical Amherstieae may be descended from a more wet-adapted subset of the African clade. The unique climatic space of the more dry-adapted lineages of Berlinia may reflect a more recent expansion into this environmental space.

### The ecological niche of the Brownea clade

The first Neotropical Amherstieae (∼56 Ma) were likely descendants of tropical African species (Bruneau et al., 2008; de la Estrella et al., 2017, 2018; Romero et al., 2020). The Brownea clade diverged in the Eocene (∼34 Ma) and diversified between 30-7 Ma (de la Estrella et al., 2017; Schley et al., 2018), a period marked by the rise of the Andes (Gregory-Wodzicki, 2000; Horton et al., 2010; Parra et al., 2010), the transition from the Megawetland Pebas system to modern Amazon landscapes at 9-8 Ma (Hoorn et al., 2010), and the formation of the Panamanian isthmus (Coates et al., 2004; Montes et al., 2015; O’Dea et al., 2016). A burst of speciation coincided with a climatic optimum between 12-7 Ma (M. de la Estrella et al., 2017; Jaramillo et al., 2006; Steinthorsdottir et al., 2021). The uplift of the Andes triggered diversification in the Neotropics (Richardson et al. 2001, Hughes and Eastwood 2006, Schley et al. 2018) because it introduced dispersal barriers and new high-elevation and open habitats, forcing allopatric distributions, and modifying climate-environmental conditions, (Morley 2003, Antonelli and Santamarin 2011, Luebert and Weigend 2014). Many endemic Andean species are derived from Amazonian ancestral lineages (Murphy et al., 2018; Schley et al., 2018).

However, our results demonstrate that the climatic space occupied by the Brownea clade remained relatively narrow, in which members are restricted only to tropical latitudes and low altitudes and did not expand into Andean landscapes or subtropical and austral mesic forests (Figs. 1, S2, S3b,d,f,h). The overlap in the E-spaces of the Brownea and Berlinia clades suggests that the environmental niche of the Brownea clade is a subset of the environmental niche of the Berlinia clade (Figs. 2, 3, S4, S5). Ecological similarities between the Brownea clade and members of the Berlinia clade have been documented in the literature, including similar occurrences in lowland tropical rainforests and along rivers (Dataset S1, Mackinder 2005, Mackinder and Pennington 2011, de la Estrella et al. 2017, Schley et al. 2018). Brownea genera, such as *Macrolobium*, are considered hydrophilic due to their habitat preference and morphological specializations, such as floating fruits (Mackinder & Pennington, 2011; Schley et al., 2018; Ter Steege et al., 2013). Although the lineages have been allopatric separated for millions of years (de la Estrella et al., 2017, 2018; LPWG, 2017; Mackinder & Pennington, 2011; Romero et al., 2020), the similarities in their occupied climatic spaces support previous research suggesting warm and wet environmental adaptations in the Amherstieae have been conserved since the early Cenozoic (Mackinder and Pennington 2011, de la Estrella et al. 2017, 2020, Schley et al. 2018).

### The ecological niche of the Berlinia clade

The Berlinia clade has a narrow temperature distribution (∼15-30° C) (Figs. 4, S1). However, the occupied climatic space of the African clade differed significantly from that of the Brownea clade (Fig. 2). These differences resulted from the bimodal environmental distribution of the Berlinia clade (Fig. 2). One cluster of Berlinia genera occupies a climatic space defined by a combination of climatic conditions distinct from the environmental space occupied by the Brownea clade (Fig. 2, Fig. S5). This unique environmental space is drier and has more variability in seasonal and diurnal temperatures.

**Figure 4.**
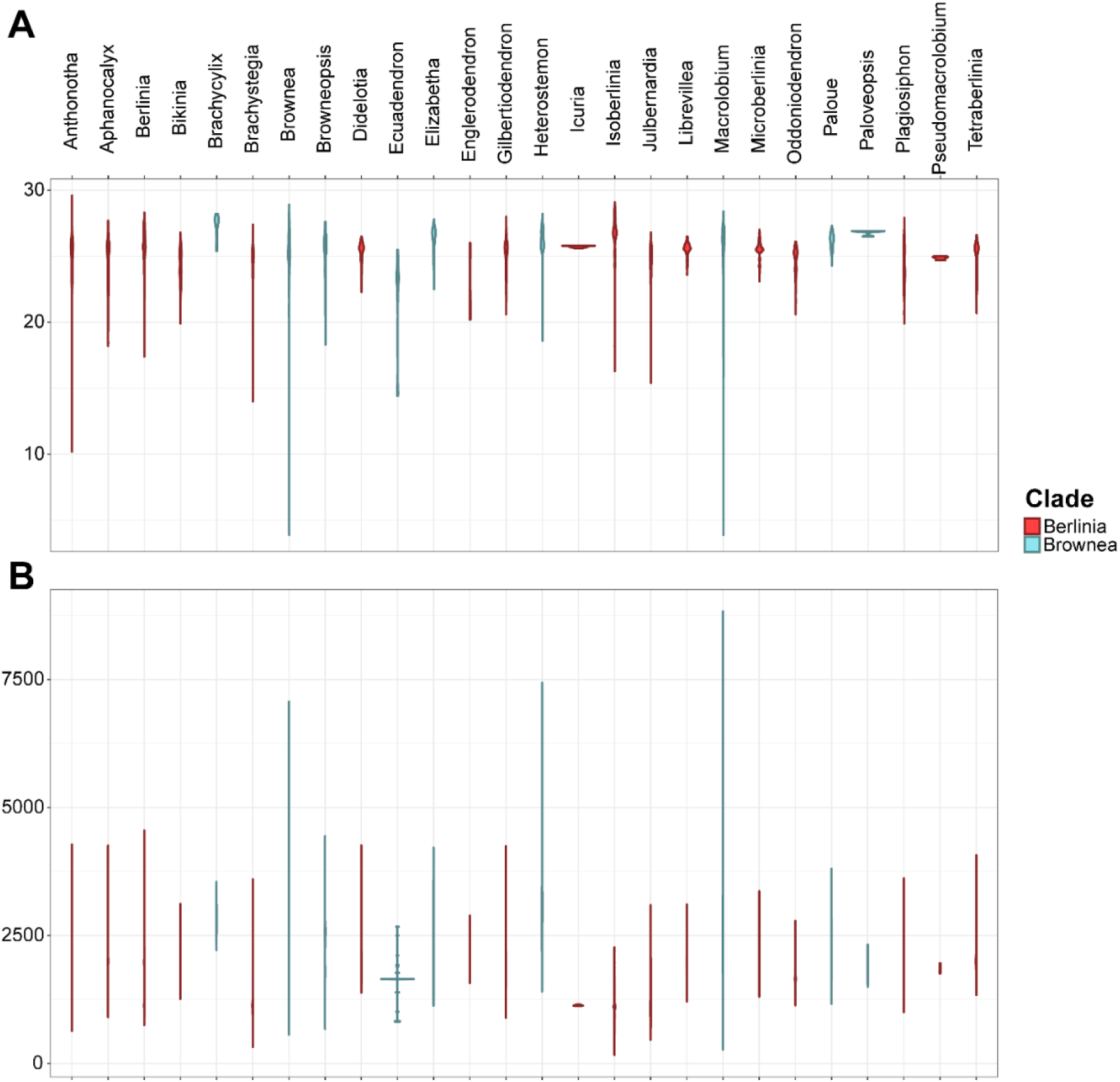
Violin density plots of mean annual temperature (°C) (A) and mean annual precipitation (mm) (B) for all genera in the Berlinia (red) and Brownea (blue) clades.

*Isoberlinia* and *Brachystegia* are two Berlinia genera distributed in drier habitats (Fig. 4). Recent phylogenetic studies time radiations within these genera to the late Cenozoic, when drier habitats expanded across Africa (de la Estrella et al. 2017, 2020). The phylogenetic ages of these lineages suggest that the expansion into this area of climate space is a more recent evolution of the Berlinia climatic niche (Wellborn and Langerhans 2015). The Berlinia clade diversified and expanded into this new drier Neogene habitat while maintaining plesiomorphic climatic adaptations in its early diverging lineages. However, the Brownea clade has not been able to occupy drier settings in South America, such as those located in Guajira (Colombia), Llanos (Colombia and Venezuela), and Cerrado (Brazil). This suggests that the common ancestor of the Brownea and Berlinia clades was adapted to wet environments with little temperature variability (Rangel et al. 2007, Wiens et al. 2010, Wellborn and Langerhans 2015). It also supports a more recent shift of the climate niche in the Berlinia clade, after the Berlinia-Brownea split in the early Cenozoic.

### Evolutionary comparisons

The higher Amherstieae diversity in the Paleotropics may reflect that Africa is the center of origin of the tribe (Fig. 1). This is also supported by the fossil record, in which the first occurrences of the tribe are registered in Africa around ∼ 58 Ma with later distributions across the tropics, ∼ 56 Ma (Romero et al., 2020). During most of the Cenozoic, African tropical rainforests were larger (Morley 2000). Tropical lineages, such as the Berlinia clade, diverged early in the Cenozoic (60-48 Ma) (de la Estrella et al. 2020). However, increased aridification in the Paleotropics during the Late Miocene reduced the extent of tropical forests and expanded savannas (Morley 2000, Plana 2004, Jacobs et al. 2010, Pokorny et al. 2015). In African Amherstieae, speciation rates potentially increased from 12 - 1 Ma while extinction rates possibly increased from ∼6 Ma and (de la Estrella et al. 2017, 2020). Most extant Berlinia genera diverged at that time (de la Estrella et al. 2020).

This shift to drier environments can be explained as an ecological opportunity that drove adaptive diversification due to niche availability (Wellborn and Langerhans 2015). In the case of the Berlinia clade, an increase in extinction rates in Amherstieae and fragmentation of tropical forests potentially opened opportunities for species of the clade to diversify and colonize warm and drier habitats (Wellborn and Langerhans 2015, de la Estrella et al. 2017, 2020). Niche shifting has been proposed as an explanation of this later diversification and adaptive response to a drier environment in Amherstieae (de la Estrella et al. 2017, 2020, Ojeda et al. 2019), in which ecological characteristics of species can change rapidly when they are established outside of their initial environmental range (Pearman et al. 2008, Guisan et al. 2014, Sherpa et al. 2009).

Another potential explanation of the Amherstieae African niche expansion is the hypothesis of unfilled niches, in which adaptive shifts could relate to niche differences induced by niche conservatism (Cunze et al. 2018, Sherpa et al. 2019). In this sense, preexisting adaptations can promote colonization of a wide range of habitats under similar climate conditions (Sherpa et al. 2019). In Amherstieae, both explanations may apply because although the subfamily Detarioideae, including the Amherstieae tribe, is predominantly distributed in tropical forests, its evolutionary history indicates the ecological flexibility to occupy both wet and dry environments due to genetic plasticity (Hughes and Eastwood 2006, Pennington et al. 2009. 2010). However, some genera in Amherstieae, such as *Isoberlinia* in the Berlinia clade, are now restricted mainly to drier habitats, suggesting a niche shift or disruptive selection (Fig 2, Fig. 4), probably resulting from multiple extinction and diversification events. This also explains the bimodal distribution of the Berlinia clade in Africa (Figs. 2, Couveur 2015, Sherpa et al. 2019, de la Estrella et al. 2020).

Conserved thermal tolerances are observed in both clades (Fig. 2, Fig. 4), similar to other legumes whose development and overall fitness are negatively affected if grown outside their preferred temperature range (Cooper 2016, Soltani et al. 2019), and are evidence of how climate stability and niche conservatism may have played a role in establishing narrow climatic ranges (Antonelli and Santamarin 2011, Wiens and Graham 2005, Pyron et al. 2015, Schley et al. 2018). Tropical forests have been climatically stable since the early Cenozoic, which explains why so many extant groups of angiosperms, including legumes, diverged in tropical regions in the early and middle Cenozoic when these environments were more extensive (Wiens 2004, Antonelli and Santamarin 2011, Pennington et al. 2010). Niche conservatism in temperature may also explain why many tropical lineages, including the Brownea clade in Amherstieae, did not expand into temperate regions or to higher elevations (Wiens and Graham 2005, Crisp et al. 2009, Pili et al. 2020).

## CONCLUSIONS

Given the ecological and economic importance of members of Amherstieae, understanding the factors that drove their distribution and how this distribution may be affected by climate change are critical questions in the conservation of the overall tribe. By comparing the climatic ranges of two clades of the Amherstieae tribe, Berlinia and Brownea, we show that the ecological niches of the African and Neotropical clades of Amherstieae demonstrate how ecological traits are conserved at taxonomic ranks higher than species, among different biogeographic regions, and despite tens of millions of years of allopatric separation. Our results suggest that the ecological niche of the Neotropical Brownea clade is a climatic subset of the African Berlinia clade, reflecting the ancestral condition of wet environment adaptations. Our results also indicated a potential expansion of the environmental niche of the African clade to include drier tropical environments as these new habitats expanded in the Neogene. However, we did not observe an expansion of the EN of the Neotropical clade to colder climates in the Andes regions, which indicates conserved thermal tolerances to warm temperatures in Amherstieae. The results of this study highlight the critical importance of conserving the tropical environments occupied by this tribe, given its ecological and economic importance.

## DATA AVAILABILITY

The data that support this study are provided in the supplementary section.

