## Supplementary material for "Niche conservatism in the Legume Amherstieae tribe: Insights from the tropical Berlinia and Brownea clades"

**Table S1.** Information on bioclimatic variables extracted from WorldClim version 2.0 (Fick and Hijmans 2017).

| ID | Acronym | Climatic variable |
|---|---|---|
| BIO1 | MAT | Annual mean temperature |
| BIO2 | MDR | Mean diurnal range (mean of monthly (max temp - min temp)) |
| BIO3 | Isothermality | Isothermality (BIO2/BIO7) (*100/0) |
| BIO4 | TS | Temperature Seasonality (Standard deviation *100) |
| BIO5 | MaxTWaMo | Max temperature of warmest month |
| BIO6 | MinTCoMo | Min temperature of coldest month |
| BIO7 | TAR | Temperature annual range (BIO5-BIO6) |
| BIO8 | MTWeQ | Mean temperature of wettest quarter |
| BIO9 | MTDQ | Mean temperature of driest quarter |
| BIO10 | MTWaQ | Mean temperature of warmest quarter |
| BIO11 | MTCoQ | Mean temperature of coldest quarter |
| BIO12 | MAP | Annual precipitation |
| BIO13 | PWeMo | Precipitation of wettest month |
| BIO14 | PDMo | Precipitation of driest month |
| BIO15 | PS | Precipitation seasonality (coefficient of variation) |
| BIO16 | PWeQ | Precipitation of wettest quarter |
| BIO17 | PDQ | Precipitation of driest quarter |
| BIO18 | PWaQ | Precipitation of warmest quarter |
| BIO19 | PCoQ | Precipitation of coldest quarter |

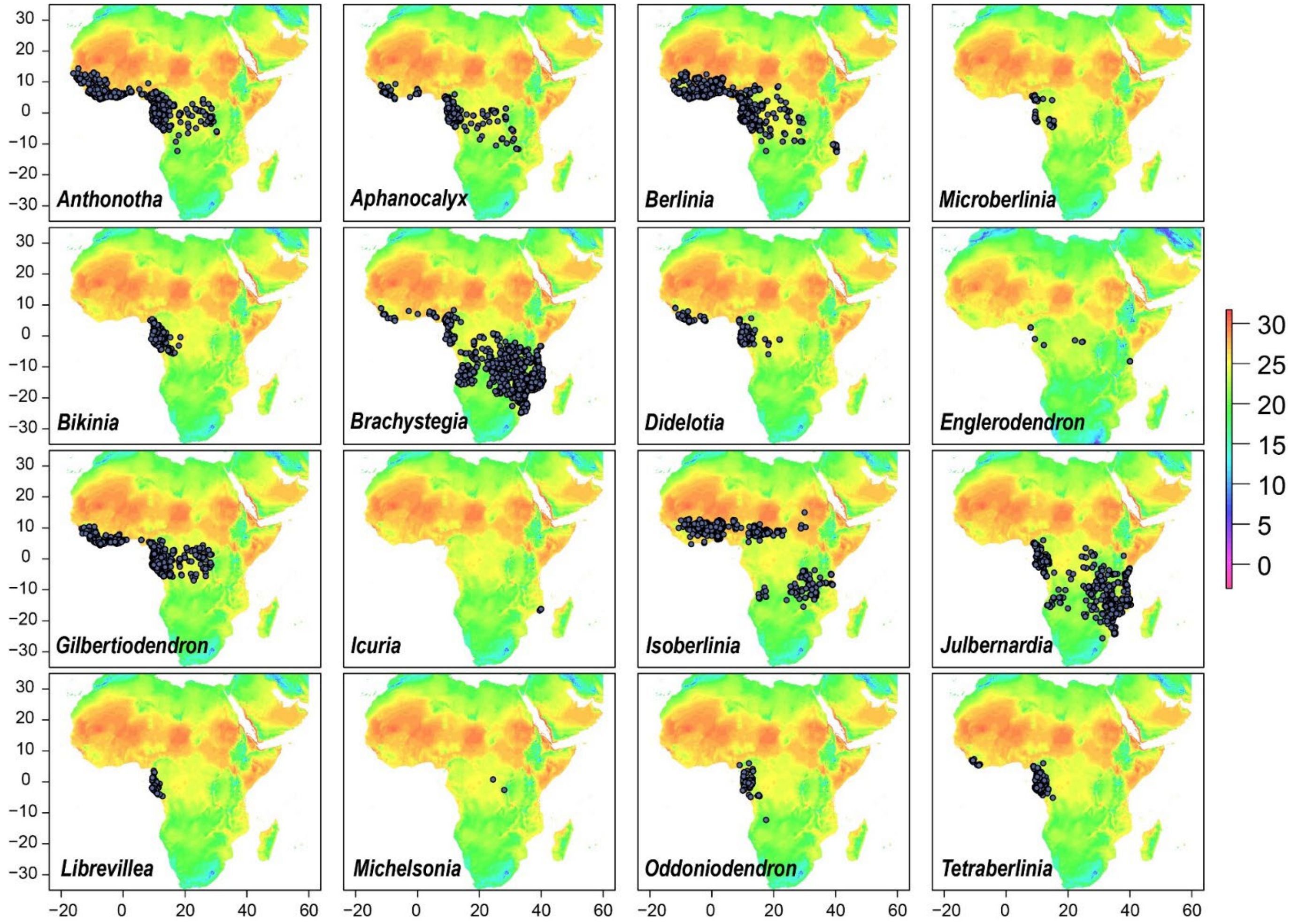


**Figure S1.** Maps show the biogeographic distribution of the 16 genera that compose the Berlinia clade, mapped against mean annual temperature (degrees ºC).

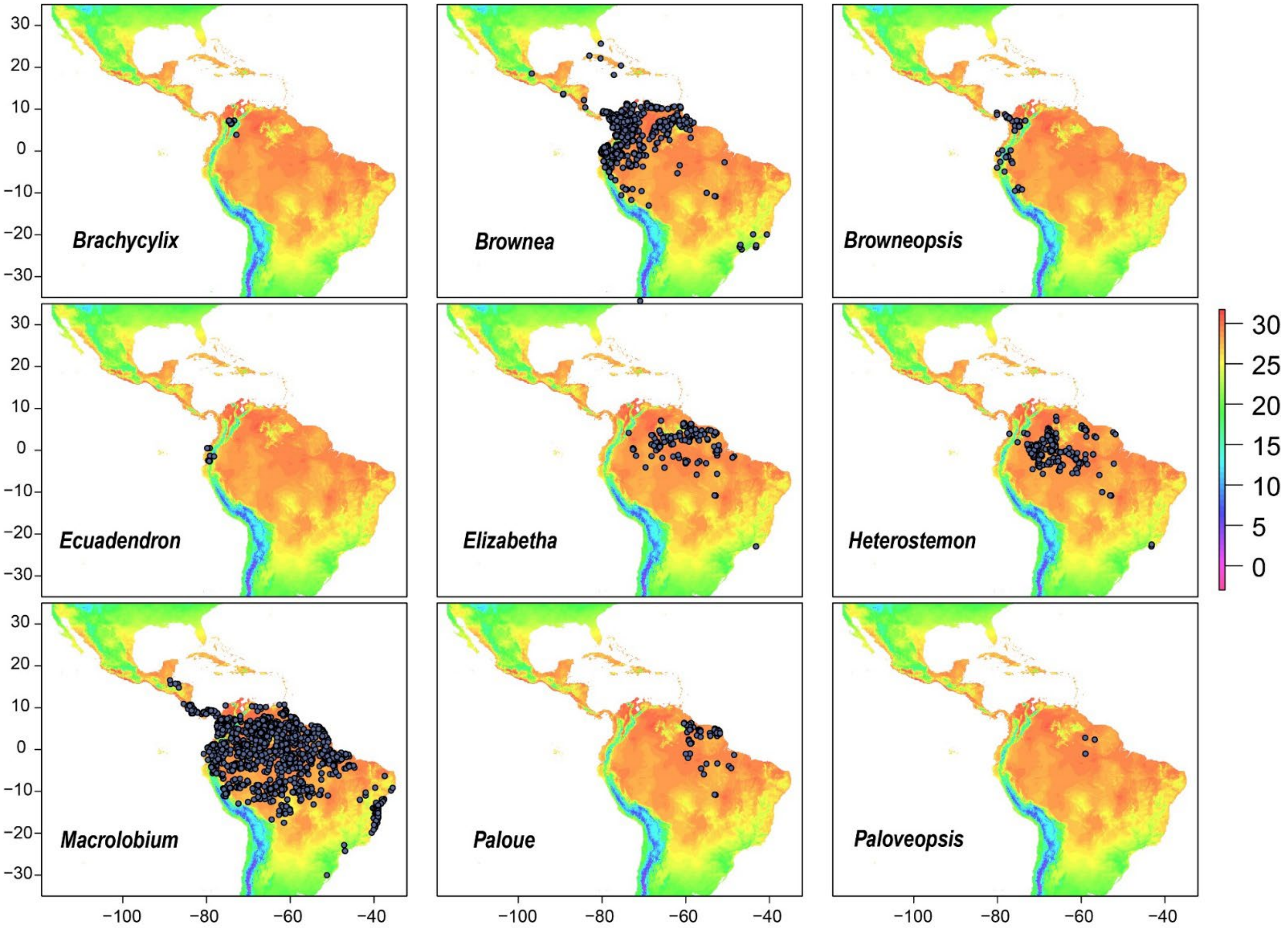


**Figure S2.** Maps show the biogeographic distribution of the nine genera that compose the Brownea clade, mapped against mean annual temperature (degrees ºC).

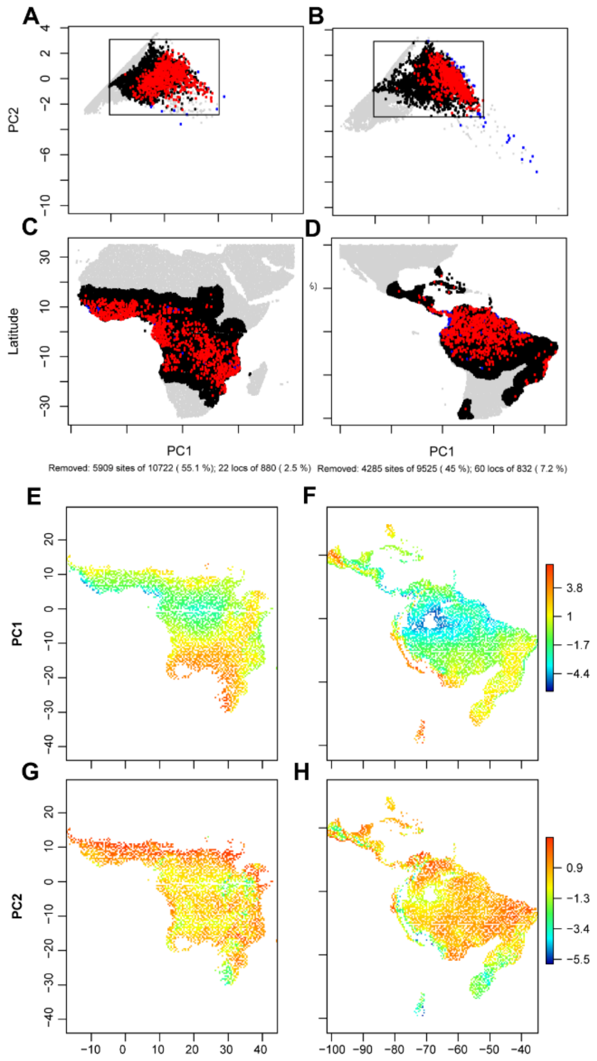


**Figure S3.** Illustrations of the analogous E-spaces for the *Berlinia* (A, C, E, G) and the *Brownea* clades (B, D, F, H). A, B) PCA plot of the environmental space for each clade. Black box depicts the shared E-space based on the maximum and minimum values of each PCs. C, D) Shared analogous E-space projected in the G-space of each clade. A-D) Black areas represent the E-space shared between both clades. Gray areas represent the non-analogous E-space of both clades. Red points indicate the occurrences of each clade retained after the E-space analysis. Blue points indicate the occurrences removed. E-H) Maps show shared accessible E-spaces for the *Berlinia* and *Brownea* clades projected into G-space. Colors correspond to PC1 (E, F) and PC2 (G, H) values.

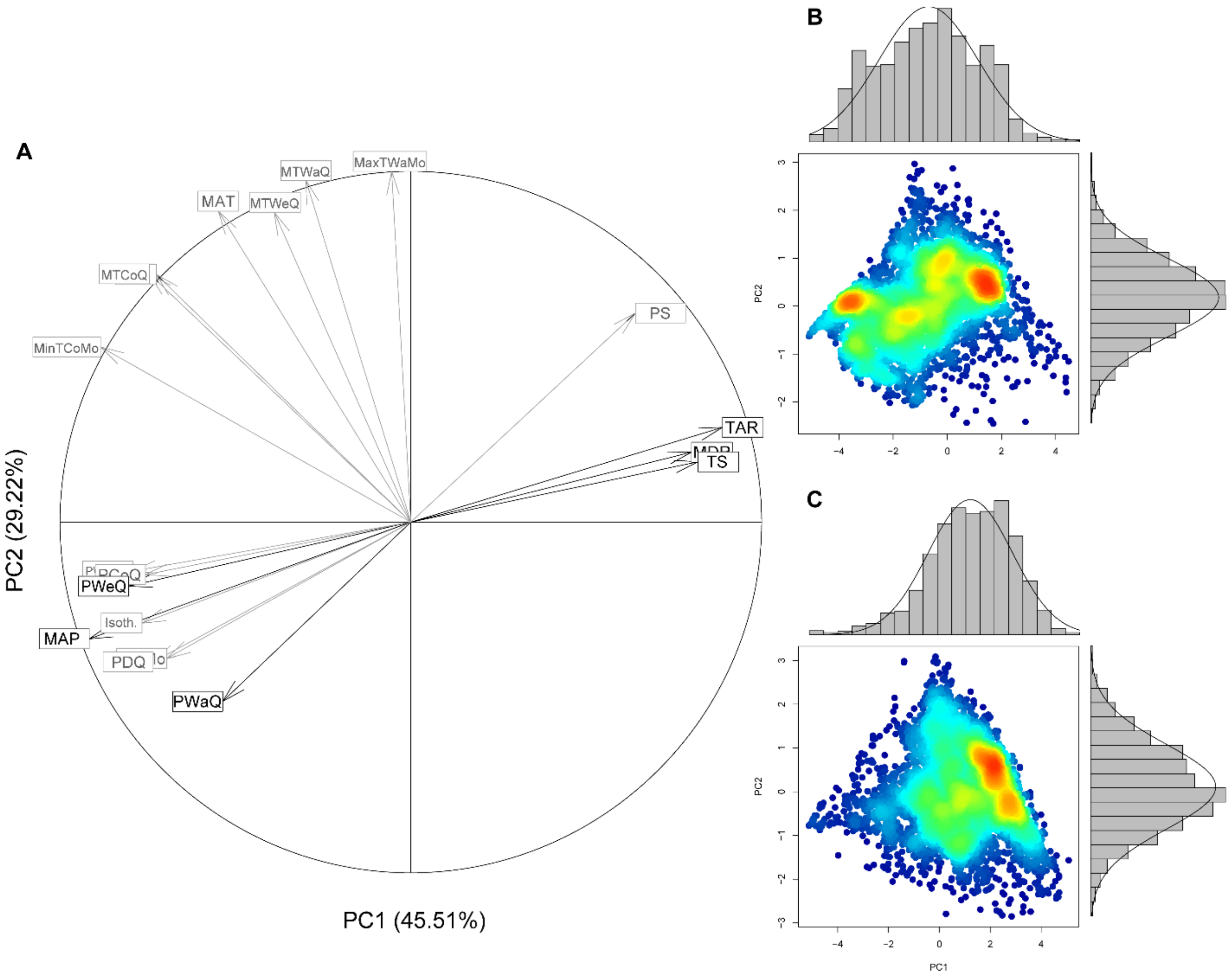


**Figure S4.** Principal component analysis of the environmental space occupied by the Berlinia and Brownea clades. A) PCA vectors illustrating the direction and magnitude of all the 19 climatic variables (see table S2). Black vectors illustrate the direction and magnitude of the six most significant environmental variables. The first two principal components explain 74.73% of the environmental variation of these clades. B, C) PCA with density plots of the occurrences and E-space for Berlinia (B) and Brownea (C) clades. Higher densities are represented with warmer colors (i.e. red) and lower densities with cooler colors (i.e. blue).

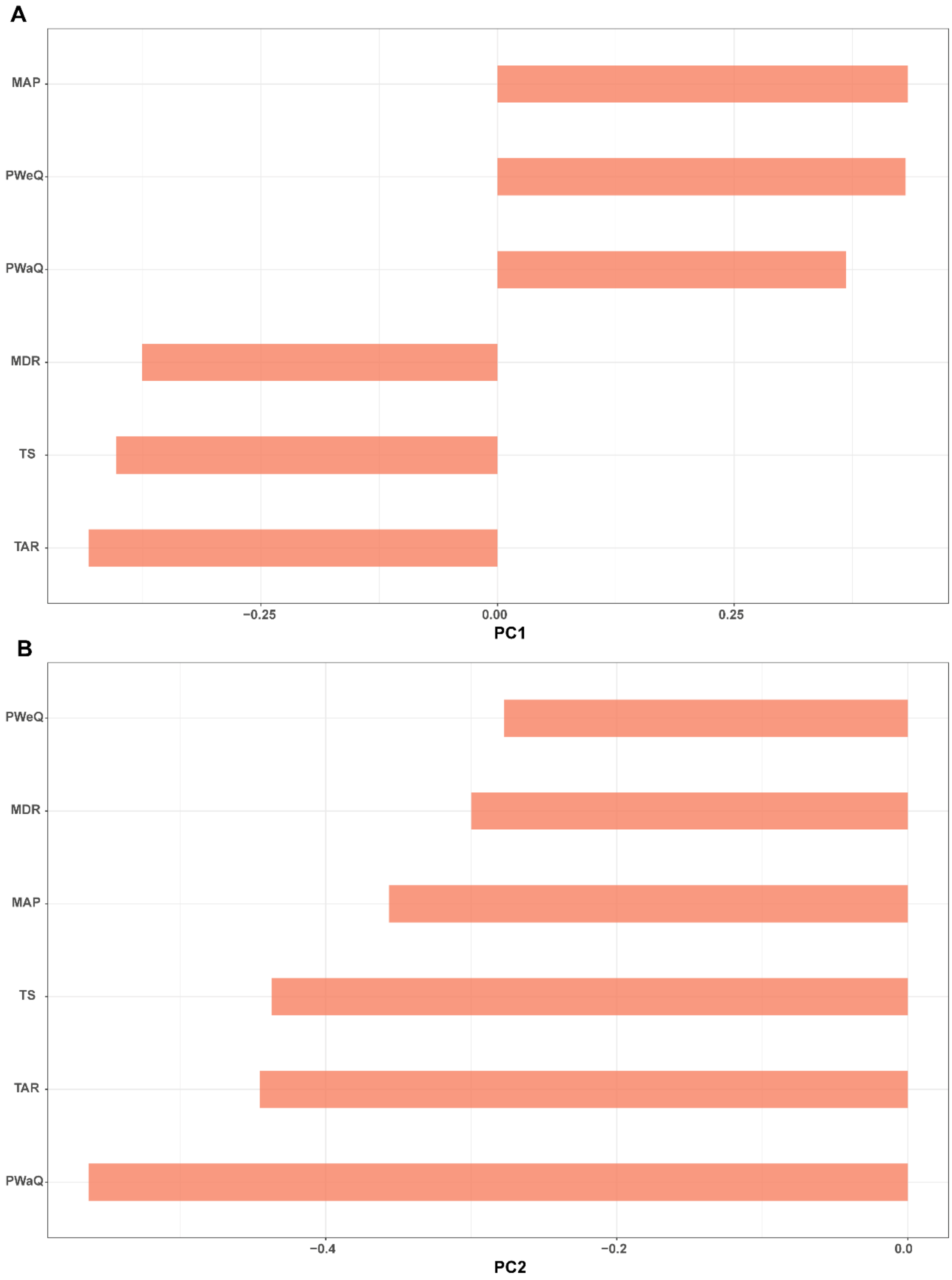


**Figure S5.** Bar plots show eigenvector loadings for PC1 (A) explaining 68.58% of the variability of the data, and PC2 (B) explaining 14.96 % of the variability of the data.
